# FIFO ATP synthase responds to glycolysis inhibition by localization into the inner boundary membrane

**DOI:** 10.1101/374967

**Authors:** K. Zalyevskiy, F. Hager, C. P. Richter, K. Psathaki, T. Appelhans, K.B. Busch

## Abstract

Mitochondrial F_1_F_0_ ATP synthase is the key enzyme to fuel the cell with essential ATP. Strong indications exist that the respiratory chain and the ATP synthase are physically separated within cristae. How static this organization is, is largely unknown. Here, we investigated the effect of substrate restriction on mitochondrial respiration and the spatio-temporal organization of ATP synthase. By superresolution microscopy, the localization and mobility of single labelled mitochondrial ATP synthase was determined in live cells. We found, that the ATP synthase under oxidative respiration displayed a clear localization and confined mobility in cristae. Trajectories of individual ATP synthase proteins show a perpendicular course to the longitudinal axis of the respective mitochondrion, exactly following the ultrastructure of cristae. When substrate for TCA cycle and respiration was limited, a significant proportion of ATP synthase localized from cristae to the inner boundary membrane, and only less mobile ATP synthase remained in cristae. These observations showing the plasticity of the spatio-temporal organisation of ATP synthase can explain why ATP synthase show interactions with proteins in distinct mitochondrial subcompartments such as inner boundary membrane, cristae junctions and cristae.

---

In normal cells, mitochondria are main suppliers for ATP by a process named oxidative phosphorylation, OXPHOS. Mitochondria are organelles with two membranes, an outer membrane (OM) and an inner membrane (IM). The inner membrane is further divided into an inner boundary membrane (IBM) that runs parallel to the OM, and cristae, extrusion of the IM into matrix. OXPHOS is performed by five membrane protein complexes in the inner mitochondrial membrane. Their joined task is to oxidize NADH/H^+^ and FADH_2_, generate a proton motive force across the inner membrane and synthesize ATP. NADH/H^+^ and FADH_2_ are products of full oxidation of pyruvate in the citrate cycle in the mitochondrial matrix. In tumour cells, pyruvate instead is largely converted to lactate, which can be measured as an extracellular acidification rate (ECAR). ATP is then mainly provided by substrate chain phosphorylation. This metabolic preference is known as the Warburg effect (1). Consequently, glycolysis inhibitors and knockdown of glycolytic enzymes are tested in diverse anti-cancer strategies (2). For example, inhibition of the glycolytic enzymes hexokinase (HK) and Fructrose-6-phosphate Kinase (F6pK) by 2-desoxyglucose (2-DG) resulted in considerable growth inhibition (3). The same study suggested that HK? plays a major role in providing substrate for respiration. Under normal conditions, cristae are the OXPHOS compartment (4) In cristae, another sub-compartmentation of the OXPHOS complexes is observed: While the respiratory complexes occupy the cristae sheet, ATP synthase lines up at the rims of cristae (5–8) (9,10) However, indications from immune EM studies and single particle tracking (SPT) exist that this distribution is not fixed (11) (12,13). Although the relative distance of F_1_F_0_ ATP synthase and respiratory oxidases is important for function (32), it is not known, whether for example, the spatio-temporal organization of the ATP synthase changes when respiration rates decrease. In addition, the many references to further interaction partners of the F_1_F_0_ ATP synthase in various sub-compartments of the IM, including compounds of the MICOS complex at cristae junctions (14, 15) or the ADP/ATP translocase and the inorganic phosphate carrier (16) (17) indicate a certain flexibility in the spatio-temporal organization of F_1_F_0_ ATP synthase.

Following, we investigated, how an inhibition of glycolysis would affect respiration, ATP synthesis, and particularly the spatio-temporal organization of F_1_F_0_ ATP synthase. The mobility and localization of membrane proteins was determined a modified version of by single particle tracking (SPT) and localization of fluorescently labeled proteins (18), which uses self-labeling tags instead of photo-switchable proteins. We recently introduced this as Tracking and Localization Microscopy (TALM) (13, 19). Mammalian cervix carcinoma cells (HeLa) were grown with quasi physiological concentration of 5.6 mM glucose as nutrient supply for several passages. Single cells showed an extended mitochondrial network (see Fig. 1A). Then, cells were incubated with an 5fold excess of the inhibitor 2-Desoxyglucose (2-DG; 30 mM) for prolonged time. With increasing incubation time, the number of individual mitochondria increased indicating a shift towards mitochondrial fragmentation. After 1.5h incubation with 2-DG, the number of individual mitochondria reached a peak and then decreased again. In parallel, the mean size of remaining networks decreased, while the overall number of (smaller) networks accordingly increased. From timepoint 2h after 2-DG addition, the mitochondrial network has started to recover (see Fig. 1B).

**Figure 1:**
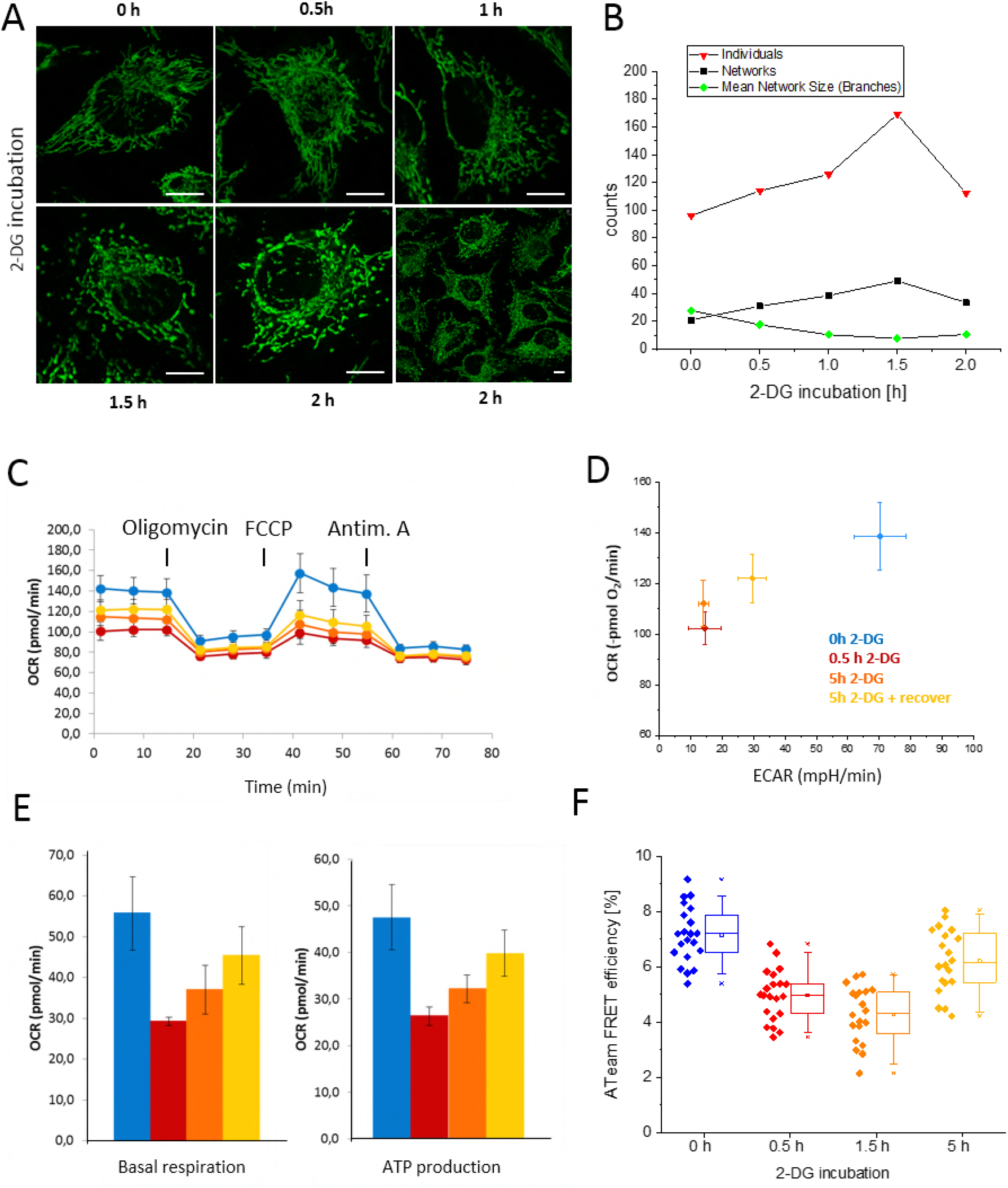
Effect of 2-DG inhibition on mitochondrial morphology and function. (*A*) Effects of 2-DG on mitochondrial morphology. (*B*) Morphometric analysis of cells incubated with 2-DG using the ImageJ plugin MiNa (20). (*C*) Mitochondrial oxygen consumption rates (OCR) at different 0.5 and 5 h after 2-DG application and 2h after recovery. Addition of inhibitor for ATP synthase (Oligomycin,1 μM), uncoupler (trifluoromethoxy carbonylcyanide phenylhydrazone; FCCP; 2 μM) and inhibitors for complex I (rotenone; 0.5 μM) and complex III (antimycin A; 0.5 μM). (*D*) Ratio of oxygen consumption rate (OCR) and extracellular acidification rate (ECAR) for metabolic phenotyping. (*E*) Basal oxygen consumption rate and ATP synthesis dependent OCR derived from (*C*). (*F*) Relative ATP levels as FRET efficiencies of an ATeam-based FRET sensor (21). Scale bars: 10 μm (A).

We next tested, whether the altered mitochondrial morphology was correlated with a different mitochondrial function. The functionality was assessed with an automatic flux analyser which allows for the determination of respiration (oxygen consumption rates, OCR) and extracellular acidification rates (ECAR) under various conditions. First, we tested respiration rates (OCR) under control conditions, after 0.5 h and 5h in the presence of 2-DG, and under conditions, where 5h incubation with 2-DG was followed by 2h recovery (see Fig. 1C). In the 2-DG samples, respiration was significantly reduced compared to the control while 2h recovery resulted in an increase of the OCR again. In order to indirectly determine ATP synthesis rates, oligomycin was added to block the ATP synthase. Under all conditions, this resulted in a drop of the OCR indicating active ATP synthesis before. Next, Trifluoromethoxy carbonylcyanide phenylhydrazonean (FCCP), an uncoupler of oxidative phosphorylation, was added to determine maximal respiration. However, while under control conditions respiration could be stimulated by uncoupling, 2-DG treated cells did not further enhance their respiration FCCP addition. Finally, respiratory dependent oxygen consumption was completely stopped by inhibiting complex III with Antimycin A, leaving only unspecific oxidation rates (see Fig. 1C). For metabolic profiling, OCR was plotted against ECAR. Already 0.5h 2-DG inhibition induced a significant drop in OCR and ECAR. 5h after addition of 2-DG, respiration was slightly increased again. When 2-DG was removed after 5h by exchange of medium with glucose for 2h, the OCR increased further and also the glycolysis block was removed indicating recovery of metabolic pathways (see Fig. 1D). In detail, 2-DG treatment reduced the basal respiration and ATP synthesis severely already after 0.5h incubation time. After 5h with 2-DG, respiration and ATP synthesis had increased again (see Fig. 1E), as confirmed by cellular ATP measurements with a FRET based sensor system (21) (see Fig. 1F). Together, these data suggest that 2-DG not only blocked glycolysis but also affected oxidative phosphorylation and mitochondrial ATP synthesis. The recovery of the respiration rates and ATP synthesis after few hours is likely due to the mobilization of a different fuel. During these time, ATP synthase went through two states: an active and an inactive state. In order to dissect, whether this was related to a specific spatio-temporal organization of F_1_F_0_ ATP synthase, we scrutinised the localization of F_1_F_0_ ATP synthase in the inner membrane with superresolution fluorescence microscopy. In addition, the mobility of single ATP synthase was determined by multiple-target tracing (13). Therefore, single, fluorescent particles were localized and tracked under the different conditions in live cells. For labeling the enzyme, subunit γ was fluorescence labeled via the self-labeling Halo7-Tag as described before (19). In the experiments, a HeLa cell line stable expressing the tag was used. As a fluorescent substrate for the self-labeling tag, Tetra-Methyl-Rhodamine-HaloTag-Ligand (TMR^HTL^) was added for 30 min (0.5-1 nM) and non-bound substrate then washed off. The low amount of TMR^HTL^ labeled particles could be distinguished as single particles. To avoid artefacts by mitochondrial crowding, F_1_F_0_ ATP synthase was imaged in the flat periphery of cells where mitochondria are separated. To achieve good signal to background ratio (SBR), a highly inclined thin illumination sheet guided through the flat part of the cells was generated with a condensing lens (22). Following, the spatio-temporal behaviour of fluorescent F_1_F_0_ ATP synthase particles was recorded in movies of 1000 to 3000 frames (50 fps). The particles were then localized by a 2D-Gaussian mask. For tracking, moving particles were traced with the multiple target-tracer (12, 23). F_1_F_0_ ATP synthase was recorded under control conditions and in the presence of the glycolysis inhibitor 2-DG for different time points and generated localization and trajectory maps (see Fig. 2A).

**Figure 2:**
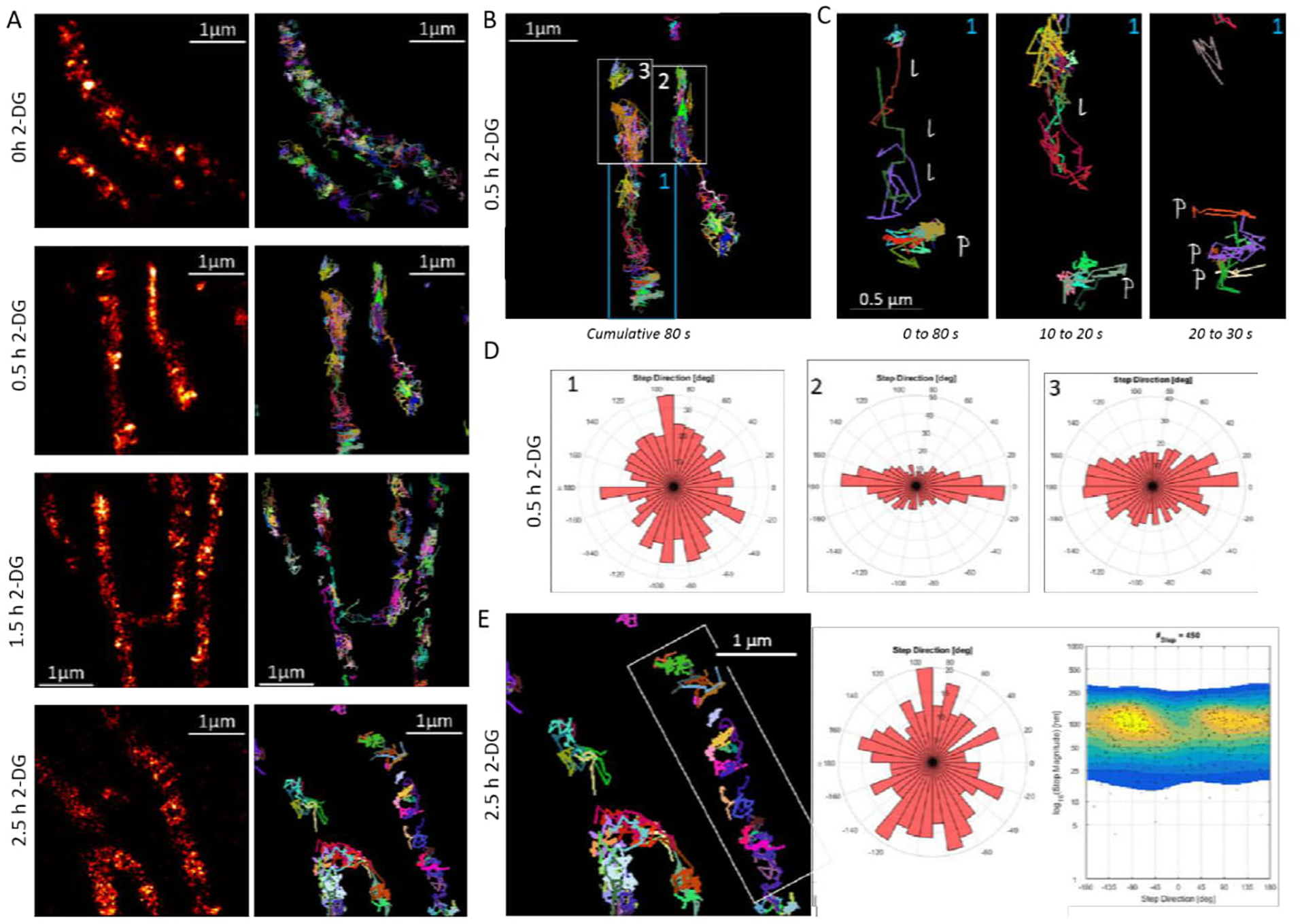
The spatio-temporal organization of ATP synthase is affected by glycolysis inhibition. (*A*) Effects of 2-DG (2-desoxyglucose) on the localization and mobility of F_1_F_0_ ATP synthase at different time points after 2-DG addition. Left panels: Localization maps of F_1_F_0_ ATP synthase in single mitochondria resulting from the summation of up to 3000 images with Gaussian’ localized F_1_F_0_ ATP synthase. Recording rate: 20 fps (50 Hz). Right panel: Trajectory maps of F_1_F_0_ ATP synthase resulting from the same data after multi-target tracing, they are also cumulative images. Localizations are depicted in *hot*. Every single trajectory in one color. (*B*) Trajectory map of F_1_F_0_ ATP synthase in two mitochondria in cells after 0.5h incubation with 2-DG. A cumulative image of trajectories of 1605 frames is shown. (*C*) Detailed view of single trajectories from the framed part 1 from (*B*), cumulative images from 500 subsequent frames (10 s) are shown. *l*: significant movement along the longitudinal axis of the mitochondrion, p: preferential movement of F_1_F_0_ ATP synthase perpendicular to the longitudinal axis of the mitochondrion. Single trajectories are differently colored; only trajectories with a minimum duration of 200 ms are shown. (*D*) The directionality of trajectories was determined for 60 ms time windows and is plotted relative to the longitudinal axis (−180°, 0°, +180°) expressed in degree. Trajectories with a directionality between −90° and +90° are assigned perpendicular; trajectories between −180° and +180° as to be longitudinal. (*E*) Mobility pattern of F_1_F_0_ ATP synthase 2.5h after 2-DG addition. Left panel: Trajectories in two different mitochondria, the framed mitochondrion is further analyzed with respect to the directionality of the trajectories. Middle panel: Step directionality (3 steps binned) of F1FO ATP synthase in the framed mitochondrion. The movement is preponderantly perpendicular, as the 360° disc and the heat map (right panel) shows. Here, 450 steps were analyzed.

The localization map of F_1_F_0_ ATP synthase in mitochondria in glucose supplied cells show distinct localization patterns and restricted diffusion due to a preponderant movement of F_1_F_0_ ATP synthase in cristae as described before (24) (see Fig. 2A, upper panel). In cells incubate for 0.5h with 2-DG glucose, the spatio-temporal map significantly changed and the arrangement of F_1_F_0_ ATP synthase was less localized but more diffuse. This was reflected in the trajectories map (0-80 s), where a conspicuous number of random trajectories appeared (see Fig. 2B). These direction-less trajectories can be assigned to F_1_F_0_ ATP synthase diffusing in the inner boundary membrane (IBM). To further dissect this, the trajectory map was then dissected into three parts, which were analyzed individually. In the framed part 1, the depiction of the trajectories from the first 500 frames (0 to 10 s) reveals a significant number of steps along the longitudinal axis (indicated as *I*), but also of steps perpendicular to the longitudinal axis (indicated as *p* in the lower part) (see Fig. 2C, left panel). During the next 10 s, other F_1_F_0_ ATP synthase trajectories along the longitudinal axis popped up (see Fig. 2C, middle panel), but also F_1_F_0_ ATP synthase moving perpendicular was present. From 20s to 30s, mainly perpendicular trajectories were recorded (see Fig. 2C, right panel). Next, we quantified this mobility behavior by dissecting the step directions. The overall direction of three consecutive steps was treated as a vector and divided into a longitudinal and perpendicular portion. For the directional analysis of mitochondria, curved mitochondria were linearized using a transformation matrix after developed after marking the longitudinal axis of the respective mitochondrion (see Fig. S1). The portion of longitudinal motion was plotted from 0 to −180°, the portion of perpendicular motion from −90° to 90°. This representation enables a quantitative overview of the preferably direction of movement. The analysis of the trajectory map from (B) shows that in the framed part #1 the movement was preponderantly perpendicular, while in the framed parts #2 and #3 the movement had a significant longitudinal share. We than analyzed the mobility of F_1_F_0_ ATP synthase in cells that were already incubated with 2-DG for 2.5 h. At least two mitochondria are present in Figure 2E, from which the mobility pattern of F_1_F_0_ ATP synthase in the framed part was further analyzed. The quantification of directionality shows a strong share of perpendicular steps for F_1_F_0_ ATP synthase under these conditions, as the distribution on the 350° disc and the heat map show (see Fig.2E).

Next, we quantified the temporal behavior of ATP synthase under the same conditions with more detail. When trajectories of selected mitochondria were analyzed with respect to their preferred directionality, the plot clearly shows that F_1_F_0_ ATP synthase in cells with normal metabolic activity (high OCR/ECAR values) displays a favored movement along the perpendicular axis of mitochondria, consistent with a confined movement in cristae (see Fig. 3A, upper panel left). With increasing time after 2-DG application, and slowing down of respiration and glycolysis, the directionality changes towards a movement along the longitudinal axis, with a peak at 1.5h after 2-DG application. After that, again more trajectories with perpendicular preference appeared leading to an equal distribution pattern with no preferential directionality (see Fig. 3A, upper panel right). For reasons of statistics and to avoid a biased interpretation, we following analyzed > 15,000 trajectories of F_1_F_0_ ATP synthase from each condition (see Fig. 3A, lower panel). The perpendicular directionality of F_1_F_0_ ATP synthase found in single mitochondria under control conditions was not that prominent any more, while the increase of trajectory directionality along the longitudinal axis after 2-DG application and reduction of metabolism was significant as already indicated when single mitochondria were analyzed. Finally, after 2.5 h, the pattern again has changed with no predominant direction any more (see Fig. 3A, lower panel right). We then determined the diffusion coefficients of F_1_F_0_ ATP synthase under the different conditions. The probability density function showed no significant differences of the average diffusion coefficient under control conditions and in presence of the glycolysis inhibitor 2-DG (see Fig. 3B), but when analyzed in detail (see Fig. S2), it was revealed that the diffusion coefficient of F_1_F_0_ ATP synthase with preferential movement in the perpendicular direction (in cristae) was significantly reduced after 0.5 and 1.5h of glycolysis inhibition, while the mobility in the longitudinal direction was stable (see Fig. 3C). Since electron microscopic images of mitochondria under control and inhibitory conditions revealed no alteration of the cristae pattern (see Fig. S3, we exclude that a change of the ultrastructure was the reason for the different mobility pattern of F_1_F_0_ ATP synthase when glycolysis was inhibited.

**Figure 3:**
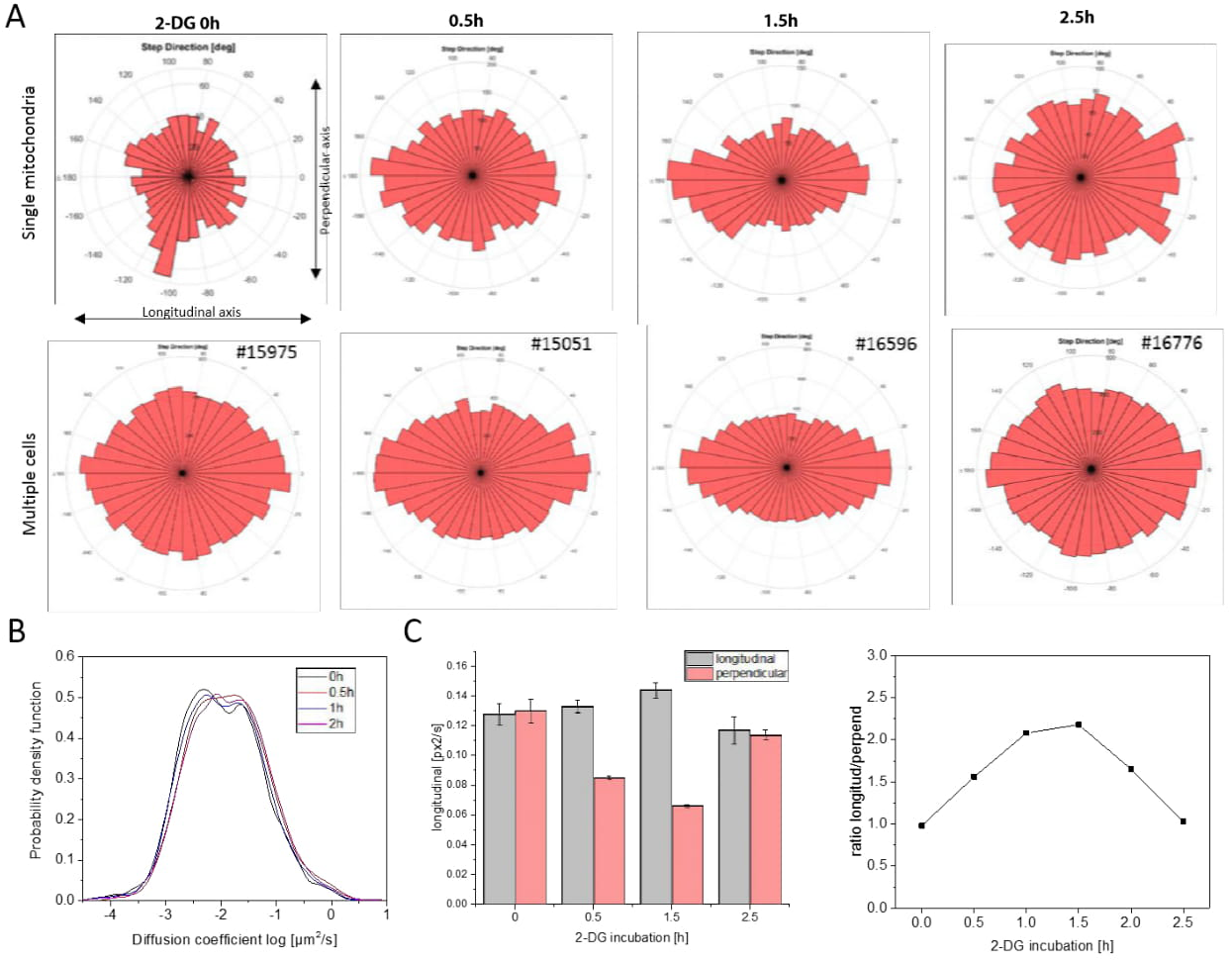
Trajectory analysis of ATP synthase under normal conditions and when glycolysis is inhibited. (*A*) Directionality of F_1_F_0_ ATP synthase trajectories in single mitochondria (upper panel). Directionality of movement obtained from > 15,000 trajectories (N>5 mitochondria). (*B*) Probability density function of diffusion coefficients under the different conditions: normal glucose (5.6 mM), 0 h; inhibition of glycolysis with 2-DG for different times: 0.5 h, 1.5h and 2.5h. (*C*) Diffusion coefficients for preferential longitudinal and perpendicular trajectories at different times after 2-DG application, and ratio of diffusion coefficients.

Together, our data suggest that (i) the spatio-temporal organization of F_1_F_0_ ATP synthase responds to an inhibition of glycolysis, (ii) this re-arrangement is characterized by an increase in the portion of ATP synthase in the IBM part, and (iii) the F_1_F_0_ ATP synthase which remained in cristae was a slow mobile fraction. The slow mobile fraction of F_1_F_0_ ATP synthase in cristae could be the molecules that are assembled as dimers and oligomers at the rims of cristae (25) (26) which keep the OXPHOS compartment in shape (5,7,27). The increase in F_1_F_0_ ATP synthase molecules that re-translocate to the inner boundary membrane would allow for interaction with proteins such as HK or VDAC, different than under conditions when F_1_F_0_ ATP synthase is coupled to respiration. In principle, a dynamic organization of F_1_F_0_ ATP synthase is prerequisite to allow for the multiple interactions of F_1_F_0_ ATP synthase, including members of the MICOS complex (14, 15) or the synthasome supercomplex with the ADP/ATP translocase and the inorganic phosphate carrier (16) (17). Also, the involvment of F_10_ ATP synthase in the formation of the mitochondrial transition pore (mPTP) (28) might require some reorganization. Our study, which shows a reorganization of the F_1_F_0_-ATP synthase in response to metabolic changes, is probably only an excerpt from the manifold commitments of ATP synthase, which make it not only plausible but necessary that the enzyme organization is dynamic.

## Acknowledgments

The authors thank Wladislaw Kohl for technical support. The study was supported by grants from the CRC SFB 944. K. Busch is associated with the CiM (cells ins motion cluster, Münster).

## Author contributions

Conception and design: K.B.B., T.A., B.R.

Acquisition of data: K.Z., B.R., F.H., K.P.

Analysis software: C.P.R.

Analysis and interpretation of data: K.Z., T.A., C.P.R., B.R., K.B.B.

Drafting or revising the article: K.B.

## Supporting Information

Supporting information is available online.

